# Machine learning aided analyses of thousands of draft genomes reveal plant- and environment-specific features of activated sludge process

**DOI:** 10.1101/710368

**Authors:** Lin Ye, Ran Mei, Wen-Tso Liu, Hongqiang Ren, Xuxiang Zhang

## Abstract

Microorganisms in activated sludge (AS) play key roles in the wastewater treatment process. However, the ecological behavior of microorganisms in AS and their differences with microorganisms in other environments have mainly been studied using 16S rRNA gene that may not truly represent their *in-situ* functions. Here, we present 2045 bacterial and archaeal metagenome-assembled genomes (MAGs) recovered from 1.35 Tb of metagenomic sequencing data generated from 114 AS samples of 23 full-scale wastewater treatment plants (WWTPs). The average completeness and contamination of the MAGs are 82.0% and 2.0%, respectively. We find that the AS MAGs have obviously plant-specific features and few proteins are shared by different WWTPs, especially for WWTPs located in geographically distant areas. Despite the differences, specific functional traits (e.g. functions related to aerobic metabolism, nutrient sensing/acquisition, biofilm formation, etc.) of AS MAGs could be identified by a machine learning approach, and based on these traits, AS MAGs could be differentiated from MAGs of other environments with an accuracy of 96.6%. Our work provides valuable genome resources for future investigation of the AS microbiome and also introduces a novel approach to understand the microbial ecology in different ecosystems.

## Introduction

Activated sludge (AS) is the largest biotechnology application in the world and is of eminent importance for the remediation of anthropogenic wastewater (Wu et al. 2019). The removal of carbon, nitrogen, phosphorus, and micropollutants is achieved by microorganisms with diverse community structures, among which populations with important metabolic functions have been individually studied (Guo et al. 2019, Kitzinger et al. 2018, McIlroy et al. 2018). Meanwhile, AS is a unique engineered ecosystem that could be controlled by a variety of operation conditions, whose attributes make it attractive for microbial ecologists to study the behaviors of microbial community assembly (Ayarza and Erijman 2011, Griffin and Wells 2017).

In the past decades, the rapid proliferation of DNA-sequencing capacity, especially high-throughput sequencing of 16S rRNA genes, boosted the understanding of the microbial community in AS (Zhou et al. 2015). One major topic is to investigate the core AS populations that are consistent occupants in a large number of parallel communities and are potentially important contributors to the system performance. Such analysis has been performed based on 16S rRNA gene sequencing in different scales, including one full-scale wastewater treatment plants (WWTPs) in Hong Kong (Ju and Zhang 2015), 13 WWTPs in Demark (Saunders et al. 2016), 14 WWTPs in Asia and North America (Zhang et al. 2012), and 269 WWTPs in 23 contries (Wu et al. 2019). Core AS microbial communities were identified at both regional and globlal scales by counting shared species or operational taxonomic units (OTUs), implying a small number of key microorganisms consititute an indispensible portion of the AS community regardless of geographical and operational variations. However, 16S rRNA gene, a useful biomarker to track evolution and construct phylogeny, does not necessarily reflect physiology and *in-situ* function (Tringe and Hugenholtz 2008). Moreover, vast metabolic diversity can still be embedded in one species or OTU, which is ususally defiened at 97% sequence identity or even higher (Yarza et al. 2014). Thus further investigation of the AS community is warrented using more advanced approaches that could resolve metabolic potentials with higher resolution.

Metagenomics aiming at recovering population genomes and annotating genetic potentials has been applied to AS and uncovered individual microorganisms or functions that are challenging to study using other methods (Albertsen et al. 2013, Perez et al. 2019, Sun et al. 2019), demonstrating that this approach is promising to reveal greater diversity at functional level than the analysis of 16S rRNA gene sequences. However, few efforts have been attempted to resolve the microbial ecology such as the core-community phenomenon in AS using metagenomics. Meanwhile, vast diversity observed in AS can present new challenges in data analysis to conventional approaches that mainly rely on reducing dimensionality, including analyses based on ordination or phylogeny of selected conserved genes. In recent years, machine-learning approaches have received growing attention and have been applied in genomics research (Eraslan et al. 2019, Zou et al. 2019). It unlike conventional methods can automatically detect patterns in data with less expert handcrafting, and therefore are suitable to handle and analyze large and complex datasets like genomics and metagenomics data (Eraslan et al. 2019, Liu et al. 2011). It can further be used to disentangle the complexity and diversity in AS community by comparing different AS systems and comparing AS with other environments.

Here, we present 2045 high- and medium-quality bacterial and archaeal metagenome-assembled genomes MAGs recovered from 114 global municipal AS samples representing one of the largest assembly of MAGs from municipal AS microbiome. After the recovery of the vast genomic information, we aimed to address two questions. First, is there a significant core AS community at MAG and protein level shared by a large number of WWTPs? Or, are there obvious plant-specific features in the AS MAGs? Second, are the AS MAGs similar to populations from other environments, Or, do they have unique environment-specific traits? In addition to a novel machine-learning approach, a collection of conventional methods analyses including genome and protein comparison, phylogenetics, ordination was applied and compared.

## Results

### 2045 MAGs obtained from AS of different WWTPs

Approximately 1.35 Tb of metagenomic sequencing data generated from 114 AS samples of 23 municipal WWTPs located in eight countries were used to construct MAGs (Figure S1, Table S1, Table S2). Among the 7548 bacterial and archaeal MAGs obtained, 2045 are estimated to have overall quality (defined as completeness - 5 × contamination) ≥ 50 (Parks et al. 2017). The average completeness and contamination of the 2045 MAGs were 82.0% and 2.0%, respectively. Figure 1A shows that 743 of the 2045 MAGs are near-complete (completeness ≥ 90, average contamination: 2.6%). The other two groups contain 845 (70 ≤ completeness < 90) and 456 MAGs (50 ≤ completeness < 70) and their average contamination values are 3.3% and 0.92%, respectively. The average contig number of these MAGs is 292, and the contig numbers have a moderate association with contamination level (Spearman’s rho = 0.47, P < 2.2e-16) but not with completeness level (Spearman’s rho = −0.11, P = 4.3e-08) (Figure S2).

**Figure 1.**
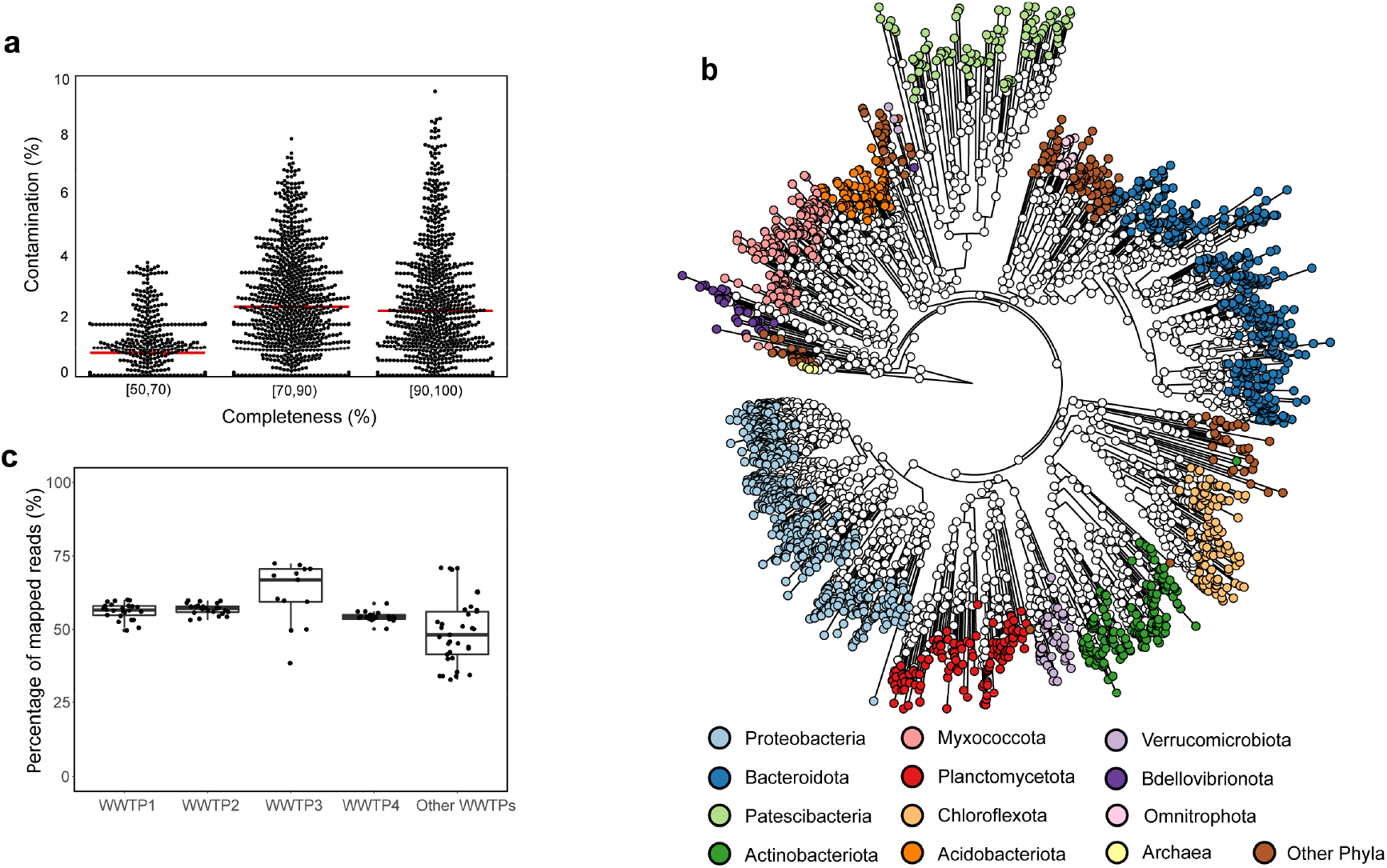
Overview of the 2045 MAGs assembled from 114 AS microbiomes. (a) Estimated completeness and contamination of the 2045 MAGs. The position of each horizontal red line refers to the mean contamination value of the corresponding group. (b) A maximum likelihood phylogenetic tree of the AS archaeal and bacterial MAGs based on universal core gene markers. The genome phylogenetic tree was generated using the universal PhyloPhlAn markers conserved across the bacterial domain. 98 MAGs with less than 80 universal markers were not included in this tree. The taxonomy of the MAGs was determined using GTDB-Tk and shown in different colors. (c) Percentages of metagenomic sequencing reads of the different AS samples mapped to the 2045 MAGs.

The 2045 MAGs were classified to 49 phyla (Figure 1b and Table S3). Among them, 21 MAGs were assigned to three archaeal phyla (*Halobacterota*, *Micrarchaeota* and *Nanoarchaeota*). For bacteria, the phylum containing highest number of MAGs was *Proteobacteria* (508 MAGs), followed by *Bacteroidota* (409 MAGs), *Patescibacteria* (178 MAGs), *Myxococcota* (164 MAGs), *Actinobacteriota* (161 MAGs), *Planctomycetota* (122 MAGs), *Chloroflexota* (114 MAGs), and *Acidobacteriota* (96 MAGs). The remaining MAGs were assigned to other miscellaneous bacterial phyla (Table S3). To further understand the diversity among these MAGs, phylogenetic analysis was performed using the universal core gene markers predicted from each MAG (Segata et al. 2013). Figure 1B shows that the clustering patterns in the tree are highly consistent with the taxonomy assignments, with *Proteobacteria* and *Bacteroidales* as the two most dominant clusters.

To estimate the representativeness of the MAGs for AS microbial genetic information, we mapped the metagenomic sequencing reads of each plant to the MAGs and calculated the percentage of mapped reads in each sample. As shown in Figure 1c, 54-63% reads (average by WWTP) of AS samples from the first four WWTPs, which have larger sequencing data volumes and significantly contribute to the AS MAG catalog, were mapped to the MAGs. For other WWTPs, the mapping ratios ranged from 34% to 72%.

### The AS MAGs show obvious plant-specific features

To evaluate the plant-specific features of the MAGs, we first analyzed the distribution of reads mapped to the MAGs obtained from different plants. As shown in Figure 2A, most (60-87%) of the mapped metagenomic reads from each WWTP were mapped to its own MAGs. A relatively small fraction of reads in each WWTP (approximately 33% in WWTP1, 32% in WWTP2, 35% in WWTP3 and 13% in WWTP4) were mapped to MAGs from other WWTPs. MAGs of WWTP1 and WWTPs share more mapped reads than other WWTP pairs (approximately 20% of sequencing reads of WWTP1 and WWTP2 are mapped to each other’s MAGs), likely because they are located in the same city.

**Figure 2.**
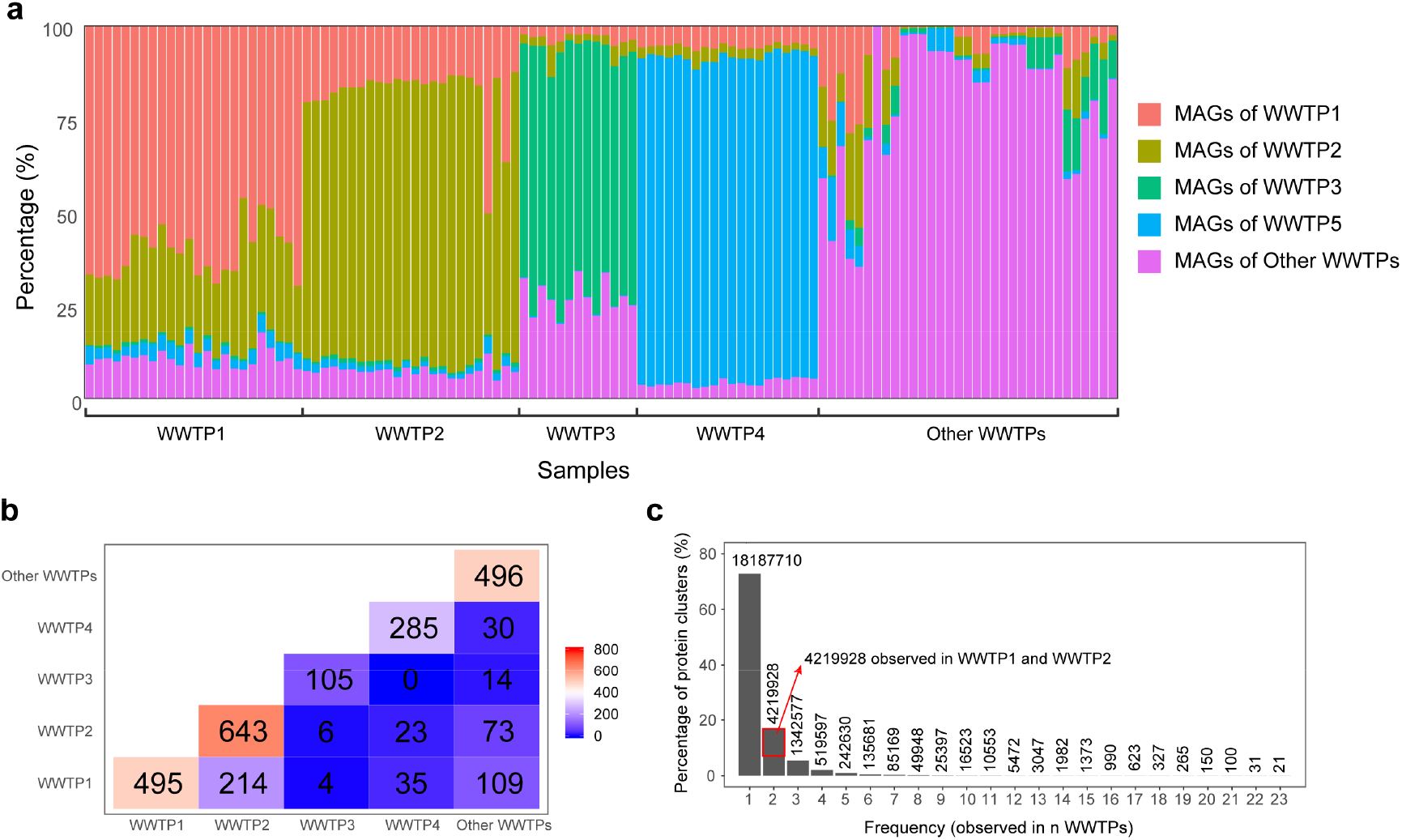
Comparison of MAGs and protein sequences among different WWTPs. (a) Relative abundance of metagenomic sequencing reads of each sample mapped to the MAGs from different WWTPs. (b) Numbers of MAG pairs with ANI>95% between different WWTPs. The values on the diagonal also refer to the MAG number in each of the first four WWTP and the total MAG number of other WWTPs (c) Frequency distribution of protein clusters across WWTPs. The protein sequences predicted from all assembly contigs were clustered at an identity cut-off of 90% with CD-HIT, then the protein clusters observed at each frequency were counted. The y-axis values were transformed to percentages and the numbers on the top of bars refer to the absolute values of protein clusters observed in n WWTPs.

Besides mapping reads to MAGs, we also computed the average nucleotide identity (ANI) values by comparing the MAGs with an all-against-all strategy. The results in Figure 2B showed that 214 MAG pairs have ANI>95% between WWTP1 and WWTP2, suggesting that these 214 bacterial or archaeal species (43% MAGs in WWTP1 and 33% MAGs in WWTP2) were shared between WWTP1 and WWTP2. However, the numbers of potentially shared species between other WWTPs were relatively small. For example, no MAG pairs with ANI>95% were observed between WWTP3 and WWTP4, and only four MAG pairs with ANI>95% were found between WWTP1 and WWTP3. A number of MAG pairs were also observed between WWTP1 and “other WWTPs” (109), and between WWTP2 and “other WWTPs” (73). This is probably because a large ratio (9/19) of WWTPs in “other WWTPs” are located in China and near to the WWTP1 and WWTP2 (Table S1).

Since the MAGs only represent part of the AS microbiome, we also conducted a pairwise comparison of protein sequences predicted from all assembled contigs of all WWTPs. As shown in Figure S3, 62% of proteins predicted from WWTP1 MAGs are highly similar (identity > 90%) to those of WWTP2 MAGs. However, only a small number of proteins predicted from WWTP3 (10-27%) and WWTP4 (7.9-28%) MAGs have highly similar hits (identity > 90%) in other WWTP MAGs. These observations could be expected as the DNA similarity among the sequences from different WWTPs could be lower due to the third-letter degeneracy. We further identified 24,850,093 clusters (identity cutoff: 90%) from the 44,212,953 protein sequences predicted from all AS samples. The frequency distribution plot (Figure 2c) shows that 73.2% of the protein clusters were found in one WWTP, and 17.0% were found in two WWTPs. Among the protein clusters observed in two WWTPs, over half (57.8%) were shared by WWTP1 and WWTP2, which were located in the same city. Only 0.1% of total protein clusters were present in ≥ 10 WWTPs. The protein comparison results confirmed the results of reads mapping and ANI calculation. It further suggested that, although a certain amount of proteins and MAGs may be shared by different WWTPs, a large proportion of bacterial populations in different WWTPs are largely different at both DNA and protein level, i.e., the bacterial genomes have plant-specific features.

### Phylogeny and functional features cannot well separate MAGs from AS and other environments

Besides comparing MAGs among different WWTPs, we also explored whether those 2024 AS MAGs obtained in this study could be distinguished from MAGs of other non-engineered (natural and animal/human-related) environments, where over seven thousand non-AS bacterial MAGs were obtained (Parks et al. 2017). We first constructed a maximum likelihood phylogenetic tree encompassing 1000 randomly-selected AS MAGs and 1000 randomly-selected non-AS MAGs (Figure 3A). The tree shows that both AS and non-AS MAGs are distributed in a wide range of phyla. The *Firmicutes* clade was dominated by non-AS MAGs (containing only 2% AS MAGs). *Myxococcota* (93% AS MAGs) and *Planctomycetota* (80% AS MAGs) have more AS MAGs than non-AS MAGs. Most of the remaining clades harbor considerable amounts of both AS and non-AS MAGs. Overall, the large-scale phylogenetic analysis based on random selection shows that the AS MAGs are phylogenetically interspersed among non-AS MAGs and no clear separation patterns could be observed.

**Figure 3.**
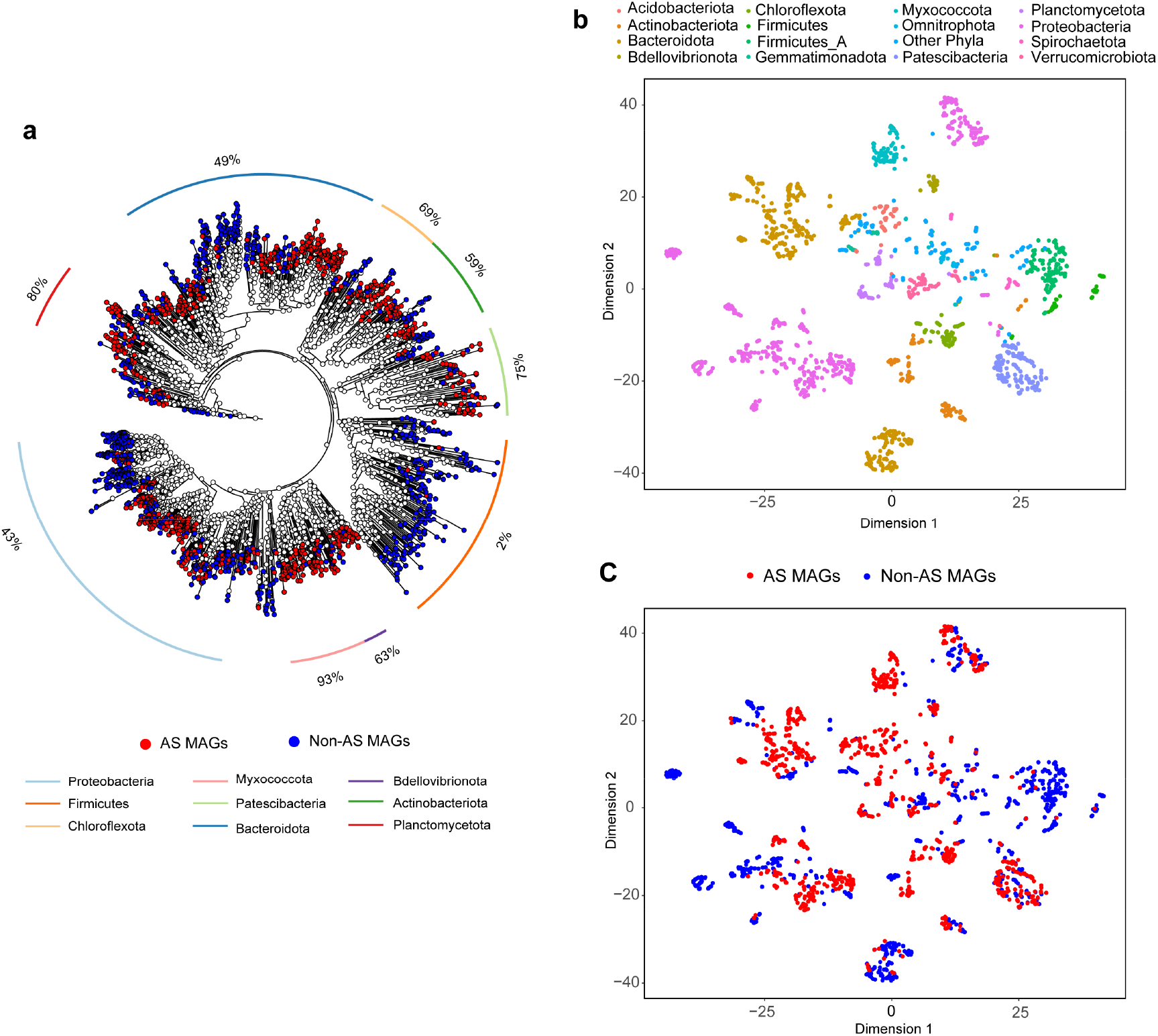
Phylogenetic and functional comparison of AS MAGs and non-AS MAGs. (a) A whole-genome maximum likelihood phylogenetic tree consisting of AS MAGs and non-AS MAGs. 1000 MAGs randomly selected from AS bacterial MAGs and 1000 MAGs randomly selected from other environments (Parks, et al. 2017) were used to build this whole-genome tree with the same methods of Figure 1b. The outside percent value refers to the relative abundance of AS MAGs in each clade. (b) Clustering of the AS and non-AS MAGs based on COG presence/absence matrix with t-SNE algorithm. The 2000 MAGs in (a) were used to generate this figure. The points representing MAGs were colored according to the taxonomy of each MAG (c) The same clustering plot with (b), while, the red points represent AS MAGs and blue points represent non-AS MAGs.

We further investigated the differences between AS and non-AS MAGs by annotating them with the COG database. As proteins in each COG have the same domain architecture and likely have the same function (Tatusov et al. 2000), comparison of COG profiles may reflect the different functions encoded in the MAGs. A COG presence/absence matrix was generated for the 2024 bacterial AS MAGs and 7164 non-AS bacterial MAGs. The t-SNE analysis based on the COG presence/absence matrix was able to separate MAGs associated with different phyla (Figure 2B). However, no clear grouping patterns between AS MAGs and non-AS MAGs (Figure 2C) was observed, which was similar to the results of the phylogenetic tree. Most of the AS and non-AS MAGs were widely distributed and co-existed in most phyla, except that few AS MAGs were observed in *Firmicutes* and some AS MAGs were separated from non-AS MAGs in the *Bacteroidota* cluster.

### A machine learning approach to distinguish between AS and non-AS MAGs based on COGs

We further explored whether machine learning can better distinguish between AS and non-AS MAGs. To do so, the COG presence/absence matrix generated from the 9188 AS and non-AS MAGs was used as an input of the random forest model (Figure 4). After the model was constructed and trained, its accuracy and applicability were further evaluated. Both the holdout method and k-fold cross-validation were applied to verify the model to avoid the over-fitting issue. For the holdout method, the dataset was divided into two partitions as testing (20%) and training (80%) sets. The number of trees is an important parameter affecting the accuracy of the random forest algorithm and should be tuned. As shown in Figure S4, after the tree number (n estimators) was increased to 200, the accuracy did not increase with the number of trees, and simultaneously other parameters (tree depth and max features) were also optimized (Figure S4). Based on the prediction results of the 20% testing data (Figure 5A), the overall prediction accuracy of the random forest model could achieve 96.6% (94% for AS and 97% for non-AS MAGs, Table S5). Particularly, the recall (i.e. true positive rate) for non-AS MAGs is 98%, which is higher than that of the AS MAGs (91%). This suggests that around 9% of AS MAGs were wrongly classified as non-AS MAGs. The F1-score, which is the harmonic average of the precision and recall, of AS and non-AS MAGs are 0.93 and 0.98, respectively. The classification accuracy obtained from 10-fold stratified cross-validation ranged from 95.0 to 95.6% (Figure 5B), suggesting that the model is reliable and accurate, and no overfitting was observed. The receiver operating characteristic (ROC) curves also demonstrated excellent performance (Area under the ROC curve (AUC) ranged from 0.94 to 1; for the mean ROC curve, AUC=0.98) of the random forest model (Figure 5C).

**Figure 4.**
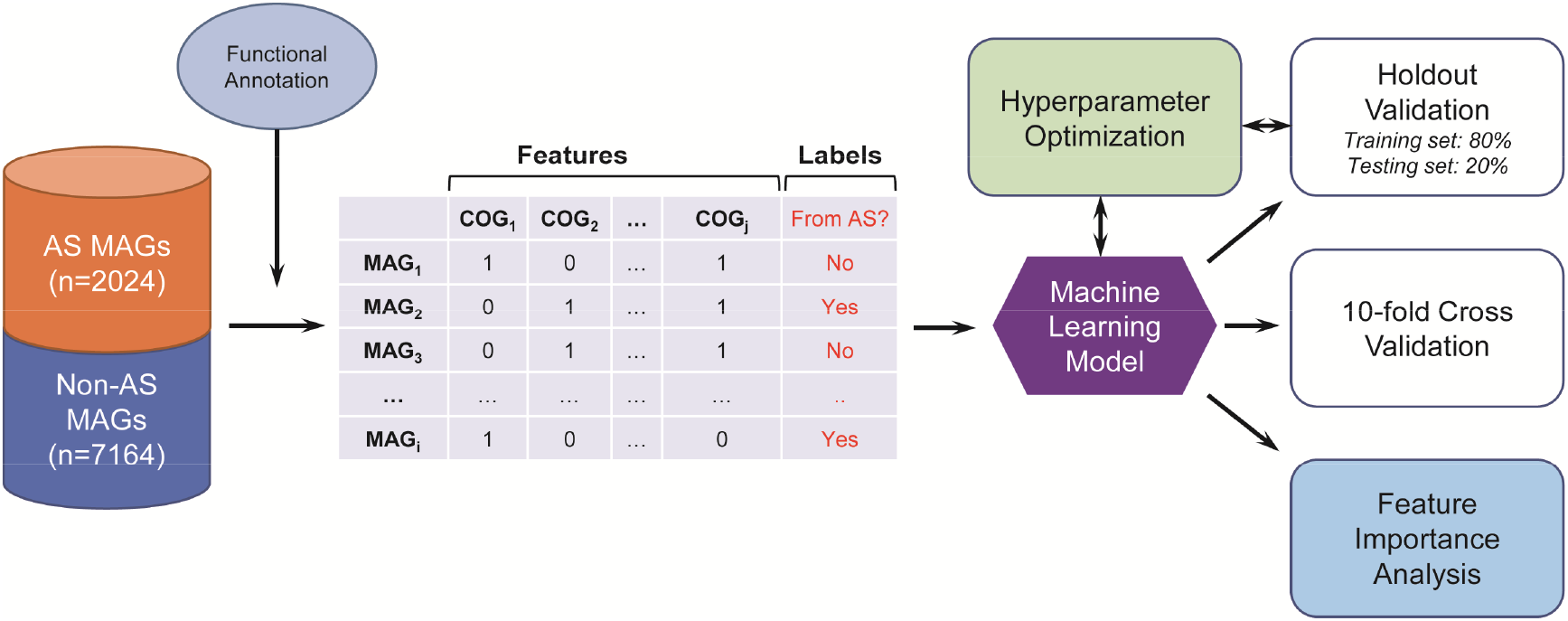
Flowchart of the implementation of machine learning for predicting AS and non-AS MAGs.

**Figure 5.**
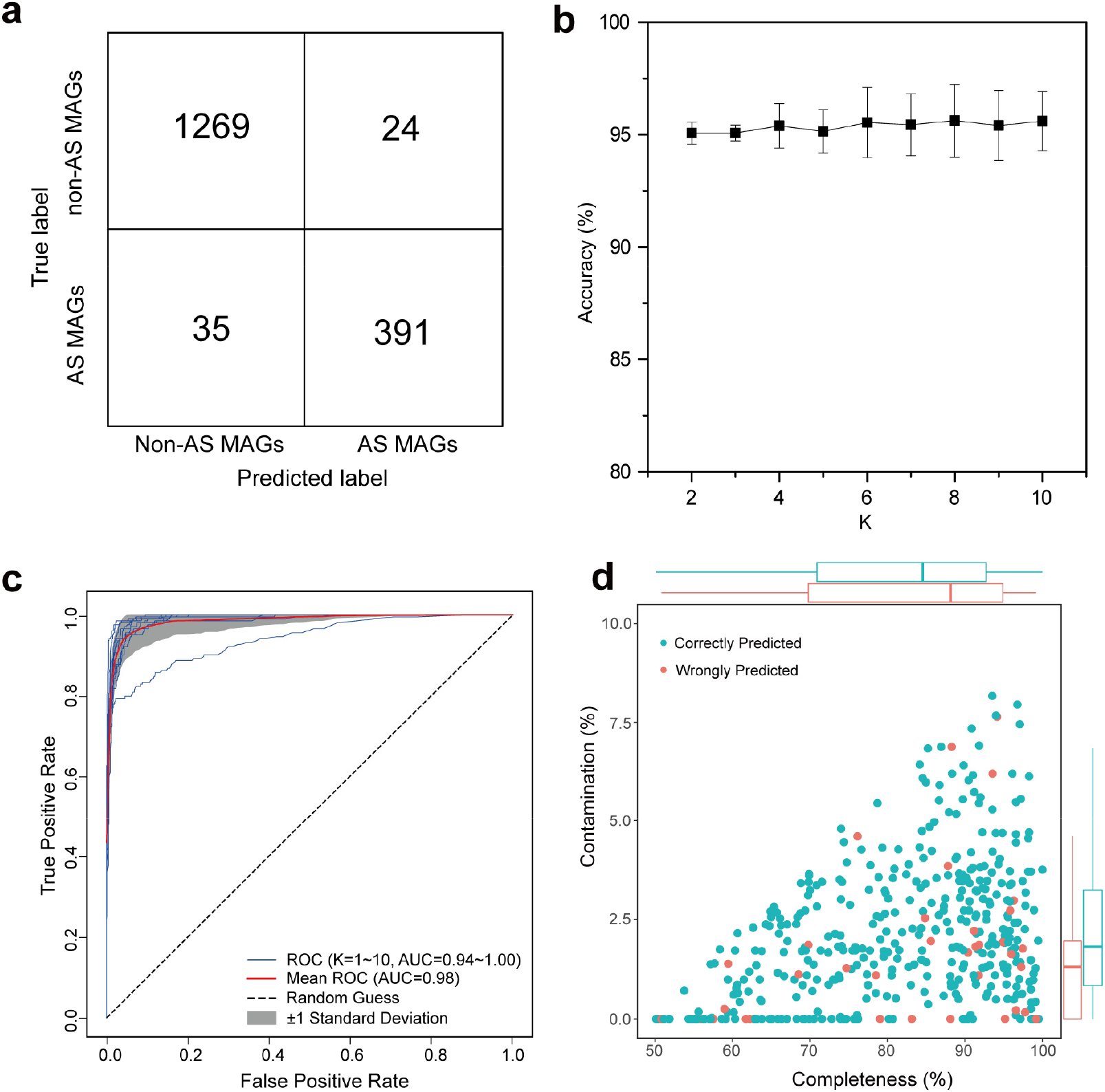
Performance of the random forest model. (a) Confusion matrix showing the performance of the random forest model on the 20% testing data of the hold-out validation. (b) Prediction accuracy of the random forest model determined based on 10-fold cross-validation. (c) ROC curves for evaluating the random forest model created from 10-fold cross-validation. (d) The completeness and contamination of correctly predicted MAGs and erroneously predicted MAGs. Boxplots along the x and y axes show the means and quartiles of the completeness and contamination values of correctly and erroneously predicted MAGs.

We further investigated the quality (completeness and contamination) and the phylogeny of the wrongly predicted MAGs. Figure 5D indicates that the wrongly predicted MAGs are evenly distributed among correctly predicted MAGs. There is no significant difference between the contamination values of the two groups of MAGs (t-test, P<0.05). The average contamination of the wrongly predicted MAGs (1.7%) is lower than that of the correctly predicted MAGs (2.2%), and the average completeness of the wrongly predicted MAGs (82.1%) is higher than that of the correctly predicted MAGs (81.6%). This suggests that the overall quality of wrongly predicted MAGs is better than that of correctly predicted MAGs. Therefore, completeness and contamination levels may not be the major reasons leading to incorrect prediction. Phylogenetic analysis showed that erroneously predicted MAGs were distributed in various phyla while a large number was associated with *Proteobacteria*, which was inherently diverse (Figure S5).

### Important functional features identified by machine learning

During the random forest model training process, an importance value was assigned to each COG. The COGs with higher importance values are more informative when the model is used to predict whether a MAG was from AS. Therefore, by analyzing the importance of each COG, the functions that differentiate the sources of MAGs can be identified. Figure 6A shows the presence/absence of the top 20 COGs based on the importance value among the MAGs (see Table 1 for the importance values and descriptions). It was clearly observed that some COGs (e.g., COG1979, 1328, 1464, 2011, 1636, etc.) are rarely present in AS MAGs. Proteins of these COGs are related to anaerobic metabolisms or functions, such as alcohol dehydrogenase, anaerobic ribonucleoside-triphosphate reductase. In contrast, several COGs (e.g. COG3324, 2114, 2107, 3303, etc.) are more frequently observed in AS MAGs than MAGs from other environments. Proteins of COG3324 and COG 2114 were related to sensing the nutritional contents of the surrounding media or other environmental signals (Lory et al. 2004). Proteins of COG 3033 are annotated as tryptophanase that catalyzes the b-elimination reaction of L-tryptophan to yield indole, ammonium and pyruvate, and the produced indole molecules may affect the biofilm formation and multidrug exporters (Yoshida et al. 2009).

**Table 1.** Importance values and descriptions of the top 20 COGs identified by random forest model to differentiate the AS and non-AS MAGs

| COG | Importance | Description |
| --- | --- | --- |
| COG0168 | 0.035 | Trk-type K<sup>+</sup> transport systems, membrane components |
| COG1115 | 0.025 | Na<sup>+</sup>/alanine symporter [Amino acid transport and metabolism] |
| COG0586 | 0.022 | Uncharacterized membrane protein DedA, SNARE-associated domain |
| COG0569 | 0.021 | Trk K<sup>+</sup> transport system, NAD-binding component [Inorganic ion transport and metabolism] |
| COG2011 | 0.015 | ABC-type methionine transport system, permease component [Amino acid transport and metabolism] |
| COG3033 | 0.014 | Tryptophanase [Amino acid transport and metabolism] |
| COG1236 | 0.011 | RNA processing exonuclease, beta-lactamase fold, Cft2 family [Translation, ribosomal structure and biogenesis] |
| COG1328 | 0.010 | Anaerobic ribonucleoside-triphosphate reductase [Nucleotide transport and metabolism] |
| COG1464 | 0.010 | ABC-type metal ion transport system, periplasmic component/surface antigen [Inorganic ion transport and metabolism] |
| COG0693 | 0.009 | Putative intracellular protease/amidase [General function prediction only] |
| COG0861 | 0.009 | Membrane protein TerC, possibly involved in tellurium resistance [Inorganic ion transport and metabolism] |
| COG1636 | 0.008 | Predicted ATPase, Adenine nucleotide alpha hydrolases (AANH) superfamily [General function prediction only] |
| COG3324 | 0.007 | Predicted enzyme related to lactoylglutathione lyase [General function prediction only] |
| COG2107 | 0.007 | Menaquinone biosynthesis enzyme MqnD [Coenzyme transport and metabolism] |
| COG1979 | 0.007 | Alcohol dehydrogenase YqhD, Fe-dependent ADH family [Energy production and conversion] |
| COG1883 | 0.007 | Na<sup>+</sup>-transporting methylmalonyl-CoA/oxaloacetate decarboxylase, beta subunit [Energy production and conversion] |
| COG2252 | 0.006 | Xanthine/uracil/vitamin C permease, AzgA family [Nucleotide transport and metabolism] |
| COG2114 | 0.006 | Adenylate cyclase, class 3 [Signal transduction mechanisms] |
| COG2846 | 0.006 | Iron-sulfur cluster repair protein YtfE, RIC family, contains ScdAN and hemerythrin domains [Posttranslational modification, protein turnover, chaperones] |
| COG1884 | 0.006 | Methylmalonyl-CoA mutase, N-terminal domain/subunit [Lipid transport and metabolism] |

**Figure 6.**
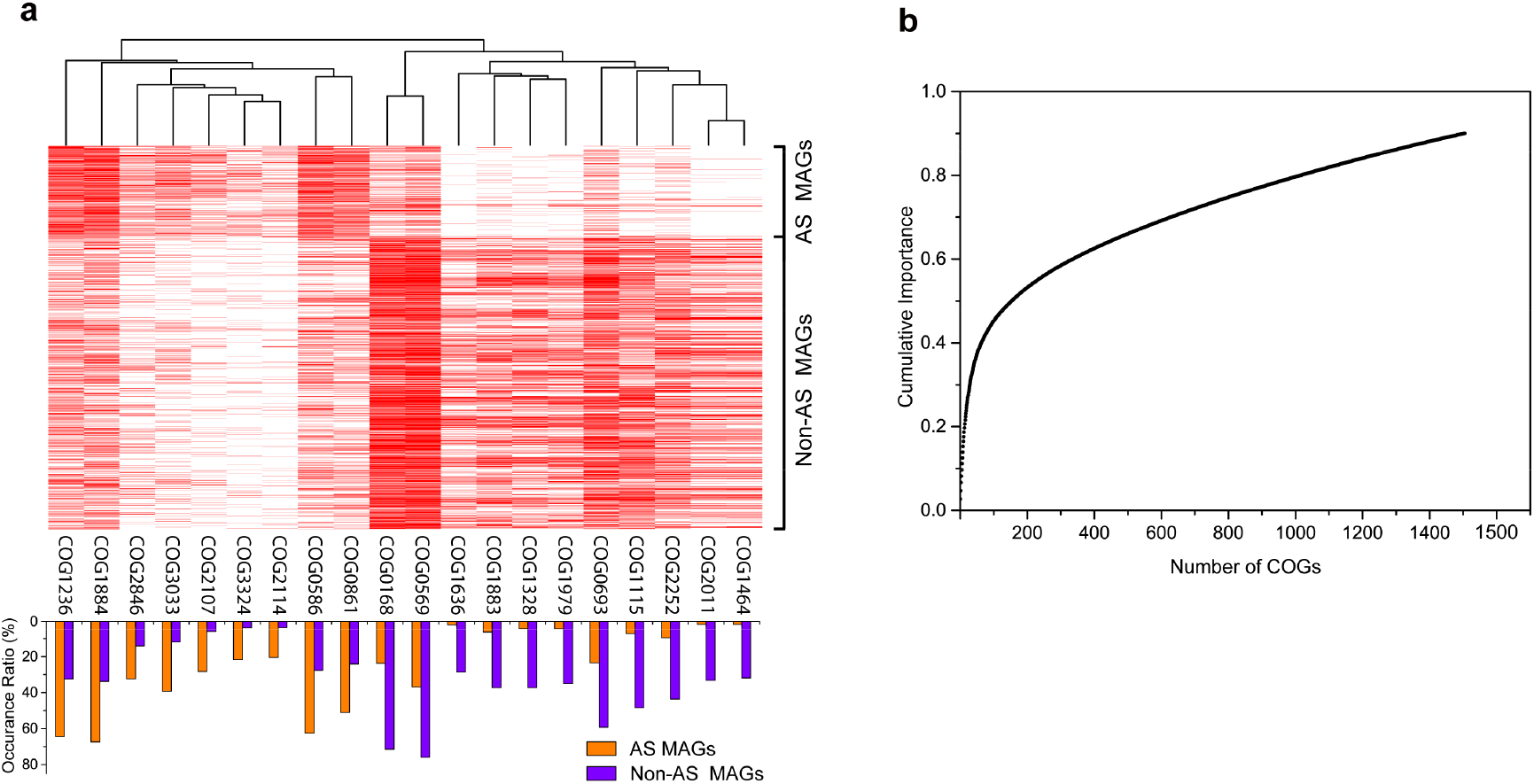
Feature importance determined by the random forest model. (a) The presence/absence of the top 20 COGs with higher importance values in each MAGs (heatmap). The bar plot shows the percentage of MAGs carrying each COG in AS MAG group and non-AS MAG group. The importance values and descriptions were shown in Table 1. (b) Cumulative importance values of the COGs.

Besides the top 20 COGs, many other COGs also contributed to the machine-learning-based prediction (Figure 6B). Among them, 148 COGs account for 50% of the cumulative importance, and approximately 1500 COGs were needed to reach a cumulative importance of 90%. This indicates the highly diverse functional features of the AS microbiomes and the strong capability of machine learning approach in capturing complex information. It also explained why the conventional phylogenetic and ordination approaches failed to separate the AS and non-AS MAGs. We found that some COGs having high importance values may associate with the aerobic condition, nutrient levels, and biofilm formation (Table 1), and they served as distinctive factors in municipal WWTP bioreactors. Meanwhile, we are still not clear with the functions of many other COGs that also significantly contribute to the machine-learning process.

## Discussion

Despite the important roles of the AS microorganisms in removing various pollutants from wastewater, the microbiome in AS remains largely uncharacterized. Based on metagenomic-assembly and binning strategies, this study constructed an AS genome catalog consisting of 2024 bacterial and 21 archaeal MAGs recovered from 114 global municipal AS samples. This catalog likely represents the largest reported AS genome collection. Its coverage of the bacteria in AS systems is considered to be high as 50-70% of the metagenomic sequencing reads could be mapped to the MAGs. Thus, it could enable us to comprehensively profile the AS bacterial community structures and functions in a higher-resolution manner.

We found that the bacterial MAGs obtained from different WWTPs could be largely different according to DNA and protein comparison, especially for WWTPs located in geographically distant areas. This suggests that AS MAGs may have plant-specific features at the genetic level, and is consistent with a recent study based on 16S rRNA gene sequencing showing that municipal AS has a small, global core bacterial community (Wu et al. 2019). Since MAGs contains much more genetic information and more variants than 16S rRNA genes, it can be inferred that the genomes of the bacteria within the small core determined based on 16S rRNA gene could also be largely different in different WWTPs. Therefore, it might be difficult to establish a “core MAG set” to include the bacterial genomes from different WWTPs. The observation of small-core populations is in line with the previously reported functional redundancy in AS ecosystems (Valentín-Vargas et al. 2012, Vuono et al. 2015). Although the overall functions of AS in all municipal WWTPs are for carbon and nutrient removal, different operational parameters and wastewater compositions may lead to significantly different microbial communities with similar functions in different WWTP. Moreover, we found that the similarity between MAGs of WWTP1 and WWTP2 located in the same city are higher than those MAGs of other WWTPs (Figure 2 and Figure S3). This is probably due to the similar wastewater compositions and environmental conditions shared by WWTP1 and WWTP2. This finding agreed with previous reports (Saunders et al. 2016, Zhang et al. 2012) that regional WWTPs were observed to have more core bacteria than global WWTPs. Overall, the low similarity of the MAGs and proteins among different WWTPs suggests that extremely high genetic diversity is present in the AS ecosystem.

Due to the extremely high genetic complexity in AS, the phylogenetic tree and COG ordination analysis failed to distinguish between AS MAG and non-AS MAGs. The major reason is that phylogenetic analysis and COG ordination are processes developed to reduce the dimensionality of multivariate data. For phylogenetic tree construction, only a limited number, usually a few hundreds, of genes coding universally conserved proteins are selected among 2000-3000 genes in a bacterial genome (Segata et al. 2013), leading to concomitant loss of genetic information. Further loss occurs during converting the sequencing data into the distance (distance-matrix methods), likelihood estimation (maximum likelihood methods) or in the process of discarding singular sites (parsimony methods) (Sourdis and Nei 1988) (Felsenstein 1981). The ordination methods (including t-SNE) also suffer from information loss due to the dimension reduction (Maaten and Hinton 2008). Although dimension reduction is important in some cases to summarize significant information from redundant high-dimensional data (Carreira-Perpinán 1997), it could miss the subtle dependencies in the datasets, for instance, the differences between AS and non-AS MAGs were not captured in this study. Here, we found that machine learning approach (random forest model) could accurately distinguish between AS MAGs and non-AS MAGs based on COG presence/absence because the random forest algorithm could take full advantages of high-dimensional data by constructing a multitude of decision trees (Breiman 2001).

The high predicting accuracy of machine learning also suggests that the municipal wastewater treatment plants can select bacteria with specific functions. Although the bacterial species in different municipal WWTPs could be different (Shchegolkova et al. 2016), they may have similar deterministic functional traits to adapt themselves in the AS system. This complements a recent study showing that the stochastic process is more important than deterministic factors on shaping the community assembly in AS based on 16S rRNA gene sequencing (Wu et al. 2019). The higher resolution of genome-level analysis reveal that AS bacterial genomes have specific functional traits despite stochastic community assembly. Furthermore, based on the random forest algorithm, we identified a number of these key functional traits (COGs), which significantly contribute to the prediction process. They are primarily related to the aerobic condition and nutrient sensing/acquisition in municipal WWTP bioreactors. However, a large number of other COGs and their co-occurrence contribution to the machine-learning model remain unexplained. Future efforts are needed to investigate the functions of the proteins assigned to these COGs.

Despite the high accuracy prediction of the random forest algorithm, we also noted that some false positive and false negative prediction results. Further analysis shows that these erroneous results are not due to the quality (completeness and contamination) of the MAGs, suggesting that the random forest model could well handle datasets with missing values (incomplete MAGs) and a certain level of noise (contaminated MAGs) (Tang and Ishwaran 2017). A small number of erroneous results is reasonable because AS is an open ecosystem, and extraneous microorganisms could be introduced into the AS through incoming raw sewage (Saunders et al. 2016) or upstream biological processes (Mei et al. 2019). In addition, the microorganisms in AS could also be easily spread to other environments via effluent discharge to receiving water body (Price et al. 2018). These stochastic propagation processes could not be captured by the machine-learning model and other technologies should be applied to identify these minor species.

Although high percentages of the metagenomic sequencing reads (50-75% for most samples) were included in the AS MAGs obtained in this study, a large number of bacterial genomes in the AS still remain unavailable due to the high complexity of the AS microbiome and micro-diversity issue, which significantly hampers genome assembly and binning (Albertsen et al. 2013, Nelson et al. 2016). We anticipate that these genomes also possess functional features similar to the MAGs obtained in this study, and future investigations based on long read sequencing (Loman and Watson 2015) or single cell sequencing (Woyke et al. 2017) are needed to confirm this hypothesis. In addition, although thousands of COGs were identified by the machine-learning model as important functional features to distinguish between AS MAGs and non-AS MAGs, only a few were known as protein clusters related to aerobic/anaerobic metabolisms, nutrient sensing/acquisition, and biofilm formation. Further investigation of these proteins would be beneficial to improve our understanding of the microbial ecology of the AS systems and provide theoretical foundation for optimizing the AS processes.

## Materials and methods

### Activate sludge sampling

In this study, 57 AS samples were collected from the aeration tanks of 11 full-scale municipal WWTPs in 8 cities of China for metagenomic sequencing (Table S1). For the two WWTPs in Nanjing City, time-series sampling was conducted every month from January 2014 to December 2015, and 24 samples were obtained from each of the two WWTPs. For other WWTPs, sampling was only conducted once in each plant during the period from April 2017 to July 2017. The detailed information of the WWTP was shown in Table S1. All sludge samples were fixed in 50% (v/v) ethanol aqueous solution and transported on ice to the laboratory for DNA extraction.

### DNA extraction and metagenomic sequencing

DNA was extracted from the AS samples using the FastDNA™ SPIN Kit for Soil (MP Biomedicals, Irvine, CA, USA) following the manufacturer’s protocol. The DNA concentration and quality were determined using a NanoDrop One spectrophotometer (Thermo Fisher Scientific, Waltham, MA, USA) and agarose gel (2%) electrophoresis. Metagenomic sequencing was conducted to obtain the entire genomic information from the sludge samples. DNA extracted from each AS sample was used for metagenomic library construction and then sequenced on an Illumina Hiseq X Ten platform (San Diego, CA, USA) with a paired-end (2 x 150) sequencing strategy.

### Collection of public activate sludge metagenomic data and metagenome-assembled genomes

Besides of the 57 AS metagenomes sequenced in this study, we also downloaded other 58 municipal AS metagenomic datasets reported in the previous studies for assembly and genome binning. The accession numbers and information of these datasets were shown in Table S1, Table S2 and Figure S1.

Moreover, a MAG catalog with over 7000 bacterial draft genomes recovered from the metagenomes different environments in a previous study (Parks et al. 2017) was used to prepare the input data for the machine-learning model. The MAGs obtained from the anaerobic digester and lab-scale wastewater treatment reactors in this catalog were excluded.

### Metagenomic assembly and contig binning

The metagenomic data were trimmed and quality-filtered using Trimmomatic v 0.32 (Bolger et al. 2014) with default parameters. Then, clean reads from all samples of each WWTP were assembled into contigs using MEGAHIT v1.1.1 (Li et al. 2015) with the following parameters: --k-min 41 --min-contig-len 1000. Then, the clean reads of each sample were mapped to the assembled contigs using Bowtie2 v 2.2.9 (Langmead and Salzberg 2012). A depth file was generated with jgi_summarize_bam_contig_depths included in MetaBAT2 (Kang et al. 2019) based on the mapping results. Then, draft genomes were recovered based on tetranucleotide frequency and contig abundance using MetaBAT2 v 2.12.1 (Kang et al. 2019). The quality of the recovered genome bins was assessed by using CheckM v 1.0.7 (Parks et al. 2015). Open reading frames were predicted in the assembled scaffolds using Prodigal v 2.6.1 (Hyatt et al. 2010) and Diamond v0.9.24.125 (Buchfink et al. 2015) were used to compare the protein sequences obtained from different WWTPs.

### Merging of compatible bins and genome refining

The ‘merge’ command of CheckM v 1.0.7 (Parks et al. 2015) was used to identify pairs of bins that could be merged according to the following criteria: (1) The completeness increased by ≥ 10% and the contamination increased by ≤ 1% when the bin pairs were merged; (2) the differences between mean GC of the bins were within 3%; (3) the mean coverage of the bins had an absolute percentage difference ≤ 25%; (4) and the bins had identical taxonomic classifications as determined by CheckM.

Genome refining was conducted with RefineM v0.0.24 (Parks et al. 2017). Briefly, contigs with a GC or tetranucleotide distance outside the 98th percentile of the expected distributions were identified and removed. Contigs were also removed if their mean coverage had an absolute percentage difference ≥ 50% when compared to the mean coverage of the bin. taxon_profile command of RefineM was used to taxonomically classify the genes comprising each bin, and contigs with divergent taxonomic classifications were removed with the ‘taxon_filter’ command of RefineM. In addition, contigs with 16S rRNA genes that appear incongruent with the taxonomic identity of each bin were also identified and removed with RefineM. Only MAGs with an overall quality ≥ 50 (defined as completeness - 5 × contamination) were used for downstream analysis. After genome refining, the genome taxonomy was assigned using GTDB-Tk v 0.2.1 (https://github.com/Ecogenomics/GTDBTk). Average nucleotide identity (ANI) between MAGs were determined using FastANI (Jain et al. 2018).

### Genome phylogenetic tree construction

The phylogenetic analyses were conducted with PhyloPhlAn using the ‘‘dev’’ branch of the repository (https://bitbucket.org/nsegata/phylophlan/overview). The genome phylogenetic tree was generated in Newick format using the 400 universal PhyloPhlAn markers conserved across the bacterial domain with the following options: ‘‘--diversity high --accurate --min_num_markers 80.’’ The final tree was reconstructed for visualization using GraPhlAn v1.1.3 (Asnicar et al. 2015).

### Functional genomic analysis

To identify protein domains in a genome, we annotate all of the MAGs using Prokka v 1.13.3 (Seemann 2014) with default parameters, and all protein domains were classified in different clusters of orthologous groups (COGs). Then, a COG matrix was derived with MAGs in rows and the presence/absence of the COGs in each MAGs as columns:

|  | COG<sub>1</sub> | COG<sub>2</sub> | ... | COG<sub>j</sub> |
| --- | --- | --- | --- | --- |
| MAG<sub>1</sub> | 0 | 1 | ... | 1 |
| MAG<sub>2</sub> | 1 | 0 | ... | 0 |
| ... | ... | ... | ... | ... |
| MAG<sub>i</sub> | 0 | 0 | ... | n<sub>ij</sub> |

Where, the matrix element n_ij_ equals 1 if MAG i encodes a protein ortholog belonging to COG_j_ and equals 0 otherwise.

The COG matrix was used to perform t-distributed stochastic neighbor embedding (t-SNE) analysis with the Rtsne package (https://cran.r-project.org/web/packages/Rtsne), and also used for the construction of the machine learning model.

### Development of machine learning model

The COG matrix constructed based on the functional annotation of the MAGs obtained in the present study and the previous study (Parks et al. 2017) was used to formulate the machine learning model to distinguish bacteria from municipal AS and those from other environments. The final dataset consists of 9288 MAGs (2024 from AS and 7164 from other environments) and 2580 COGs, which were used to train two machine learning models based on support vector machine and random forest algorithms respectively. Random forest was chosen because it has higher accuracy than support vector machine. Moreover, it is known that random forest algorithm is suitable for data sets with many features, especially where each of the features contributes little information (Breiman 2001).

The model training and evaluation was performed with scikit-learn (https://scikit-learn.org/), a Python package for machine learning. Both the holdout method and k-fold cross-validation was applied to verify the model. For the holdout method, the dataset is divided into two partitions as training (80%) and testing (20%) sets, respectively. The training set was used to train the model and the unseen testing data was used to test the predictive ability. Over-fitting is a common issue in machine learning which can occur in most models (Domingos 2012). In this study, out-of-bag (OOB) estimates were applied to avoid overfitting. In addition, 10-fold cross-validation was conducted to verify that the model is not overfitted. The dataset is randomly partitioned into 10 mutually exclusive and approximately equal subsets, and one set was kept for testing while others were used for training. This process was iterated with the 10 subsets. Furthermore, the COGs significantly contributed to the machine-learning-based prediction were analyzed based on the feature importance provided by the random forest model.

## Acknowledgements

We thank Ying Yang, Feng Guo, Jizhou Duan, Xingtao Zhang, Kailong Huang, Fuzheng Zhao, Mingyuan Sun, Wenda Tao, Yongchao Xie, Peng Liu, Zhuoyan Shen, Haohao Sun and Qingmiao Yu for assistance on the sludge sampling and sample treatment. This study has received funding from the National Natural Science Foundation of China (51608256, 51878333).

