## Supplementary Information for "Machine learning aided analyses of thousands of draft genomes reveal plant- and environment-specific features of activated sludge process"

**Table S1**. Information about the WWTPs from which the activate sludge samples used in this study were collected

| No. | WWTP ID | WWTP Name/Label | Process | Location | Latitude/Longitude | Sample Number | Data Size (G bps) | Reference |
|---|---|---|---|---|---|---|---|---|
| 1 | WWTP1 | JXZ | A/A/O | Nanjing, China | 32.033613, 118.703391 | 24 | 267.1 | This study |
| 2 | WWTP2 | DC | OD | Nanjing, China | 32.225880, 118.753075 | 24 | 331.1 | This study |
| 3 | WWTP3 | Aalborg West | NA | Aalborg, Denmark | 57.048139, 9.906371 | 13 | 274.7 | (Munck et al. 2015) |
| 4 | WWTP5 | Ulu Pandan | CAS+MBR | Singapore | 1.321595, 103.776771 | 20 | 180.8 | (Law et al. 2016) |
| 5 | WWTP6 | QS | A/A/O | Zhuhai, China | 22.251326, 113.523755 | 1 | 16.3 | This study |
| 6 | WWTP7 | HP | A/A/O | Guangzhou, China | 23.193854, 113.437769 | 1 | 17.1 | This study |
| 7 | WWTP8 | JM | A/A/O | Xiamen, China | 24.633361, 118.012453 | 1 | 18.0 | This study |
| 8 | WWTP9 | XC | A/A/O | Wuxi, China | 31.497265, 120.324492 | 1 | 16.7 | This study |
| 9 | WWTP10 | DJQ | CAST | Wuxi, China | 31.610036, 120.303888 | 1 | 16.7 | This study |
| 10 | WWTP11 | SZ1 | A/A/O | Suzhou, China | 31.345850, 120.630079 | 1 | 19.6 | This study |
| 11 | WWTP12 | SZ2 | A/A/O | Suzhou, China | 31.396898, 120.502044 | 1 | 18.2 | This study |
| 12 | WWTP13 | HBH | A-B process | Qingdao, China | 36.107309, 120.332498 | 1 | 15.5 | This study |
| 13 | WWTP14 | SY | A/O | Shenyang, China | 41.799910, 123.443920 | 1 | 15.3 | This study |
| 14 | WWTP15 | Manitowoc | NA | USA | 44.097347, -87.687685 | 3 | 10.8 | (Chu et al. 2018) |
| 15 | WWTP16 | Sheboygan | NA | USA | 43.753345, -87.723262 | 3 | 9.1 | (Chu et al. 2018) |
| 16 | WWTP17 | SwgA | CAS | Buenos Aires, Argentina | -34.604845, -58.509590 | 4 | 14.4 | (Ibarbalz et al. 2016) |
| 17 | WWTP18 | SwgB | CAS | Buenos Aires, Argentina | -34.633961, -58.403628 | 4 | 18.1 | (Ibarbalz et al. 2016) |
| 18 | WWTP19 | Kocejve | NA | Slovenia | 45.633202, 14.844261 | 3 | 18.2 | (McIlroy et al. 2016) |
| 19 | WWTP20 | AD | NA | Swiss | 46.708533, 6.905204 | 1 | 5.7 | (Ju et al. 2019) |
| 20 | WWTP21 | WT | NA | Swiss | 46.476471, 7.976473 | 1 | 5.3 | (Ju et al. 2019) |
| 21 | WWTP22 | TE | NA | Swiss | 47.276185, 7.445573 | 1 | 6.1 | (Ju et al. 2019) |
| 22 | WWTP23 | TU | NA | Swiss | 47.325399, 9.056555 | 1 | 5.4 | (Ju et al. 2019) |

| | | | | | | | | |
|---|---|---|---|---|---|---|---|---|
| 23 | WWTP24 | EGA | NA | Demark | 56.214686, 10.266368 | 3 | 52.0 | (Munck et al. 2015) |

**Abbreviations in the "Process" column**: A/O, anoxic/aerobic; A/A/O, anaerobic/anoxic/aerobic; CAS, conventional activated sludge; CAST, cyclic activated sludge technology; OD: oxidation ditch; MBR, membrane bioreactor; NA, not available.

**Table S2**. Accession numbers of the metagenomic datasets used in this study

| **WWTP ID** | **Accession numbers** |
|---|---|
| WWTP1 | Submission in progress |
| WWTP2 | Submission in progress |
| WWTP3 | ERR712369, ERR712370, ERR712371, ERR712372, ERR712373, ERR712374, ERR712375, ERR712376, ERR712377, ERR712378, ERR712379, ERR712380, ERR712381, ERR712382, ERR712383, ERR712384 |
| WWTP5 | SRR3501849, SRR3501850, SRR3501851, SRR3501852, SRR3501853, SRR3501854, SRR3501855, SRR3501856, SRR3501857, SRR3501858, SRR3501859, SRR3501861, SRR3501862, SRR3501873, SRR3501884, SRR3501885, SRR3501886, RR3501887, RR3501888, SRR3501889 |
| WWTP6 | Submission in progress |
| WWTP7 | Submission in progress |
| WWTP8 | Submission in progress |
| WWTP9 | Submission in progress |
| WWTP10 | Submission in progress |
| WWTP11 | Submission in progress |
| WWTP12 | Submission in progress |
| WWTP13 | Submission in progress |
| WWTP14 | Submission in progress |
| WWTP15 | SRR5570992, SRR5570991, SRR5571003 |
| WWTP16 | SRR5571009, SRR5571008, SRR5571011 |
| WWTP17 | SRR2107210, SRR2107211, SRR2107212, SRR2107213 |
| WWTP18 | SRR2107215, SRR2107216, SRR2107218, SRR2107219 |
| WWTP19 | ERR1076073, ERR1076074, ERR1076075 |
| WWTP20 | ERR2808645, ERR2808647 |

| | |
|---|---|
| WWTP21 | ERR2808646, ERR2808661 |
| WWTP22 | ERR2808648, ERR2808650 |
| WWTP23 | ERR2808649, ERR2808664 |
| WWTP24 | ERR712389, ERR712390, ERR712391 |

**Table S3**. Abundance of AS MAGs assigned to each phylum

| Domain | Phylum | Number of bins | Relative abundance (%) |
|---|---|---|---|
| Bacteria | Proteobacteria | 508 | 24.84 |
| | Bacteroidota | 409 | 20.00 |
| | Patescibacteria | 178 | 8.70 |
| | Myxococcota | 164 | 8.02 |
| | Actinobacteriota | 161 | 7.87 |
| | Planctomycetota | 122 | 5.97 |
| | Chloroflexota | 114 | 5.57 |
| | Acidobacteriota | 96 | 4.69 |
| | Verrucomicrobiota | 50 | 2.44 |
| | Bdellovibrionota | 34 | 1.66 |
| | Gemmatimonadota | 23 | 1.12 |
| | Nitrospirota | 18 | 0.88 |
| | Omnitrophota | 18 | 0.88 |
| | Elusimicrobiota | 17 | 0.83 |
| | Spirochaetota | 17 | 0.83 |
| | Eisenbacteria | 13 | 0.64 |
| | Armatimonadota | 11 | 0.54 |
| | Cyanobacteriota | 10 | 0.49 |
| | AABM5-125-24 | 6 | 0.29 |
| | UBP1 | 5 | 0.24 |
| | Firmicutes | 4 | 0.20 |
| | Zixibacteria | 4 | 0.20 |
| | Calditrichota | 4 | 0.20 |
| | KSB1 | 4 | 0.20 |
| | Campylobacterota | 3 | 0.15 |
| | Firmicutes_A | 3 | 0.15 |
| | Desulfobacterota | 3 | 0.15 |
| | UBA10199 | 3 | 0.15 |
| | OLB16 | 2 | 0.10 |
| | Thermotogota | 2 | 0.10 |
| | Margulisbacteria | 2 | 0.10 |
| | BRC1 | 2 | 0.10 |
| | Cloacimonadota | 2 | 0.10 |
| | UBP10 | 2 | 0.10 |
| | Nitrospirota_A | 2 | 0.10 |
| | Synergistota | 1 | 0.05 |
| | Fermentibacterota | 1 | 0.05 |
| | Bipolaricaulota | 1 | 0.05 |
| | Firmicutes_H | 1 | 0.05 |

| | | | |
|---|---|---|---|
| | UBP7 | 1 | 0.05 |
| | Fusobacteriota | 1 | 0.05 |
| | Hydrogenedentota | 1 | 0.05 |
| | Dependentiae | 1 | 0.05 |
| Archaea | Halobacterota | 5 | 0.24 |
| | Micrarchaeota | 6 | 0.29 |
| | Nanoarchaeota | 10 | 0.49 |

**Table S4**. Sources of the non-AS MAGs published by Parks et al. (2017)

| **Environmental sources** | **Number of MAGs** |
|---|---:|
| seawater | 1793 |
| subway | 1272 |
| gut | 1128 |
| rumen | 866 |
| soil | 833 |
| tailing pond | 448 |
| oil sand | 165 |
| freshwater | 164 |
| freshwater sediment | 111 |
| mine biofilm | 82 |
| crystal geyser | 48 |
| marine sediment | 26 |
| coral mucus | 25 |
| sponge | 24 |
| ground water | 22 |
| travertine | 21 |
| sand | 20 |
| cassava fermentation water | 16 |
| acidic hot spring | 15 |
| drinking water | 15 |
| bodily fluid | 13 |
| crude oil | 11 |
| fracture water | 11 |
| mud volcano | 10 |
| marine hydrothermal vent chimney | 8 |
| sand filter | 5 |
| house | 3 |
| sea rock | 3 |
| pinewood nematode | 2 |
| medieval human | 1 |
| mine tailings | 1 |
| oilsands tailings ponds | 1 |
| salt lake | 1 |

**Table S5**. Prediction report of the random forest model

| **Category** | **Precision** | **Recall** | **F1-score** |
|---|---|---|---|
| AS MAGs | 0.94 | 0.91 | 0.93 |
| Non-AS MAGs | 0.97 | 0.98 | 0.98 |

Precision=True Positive / (True Positive + False Positive)
Recall=True Positive / (True Positive + False Negative)
F1-score = 2*Precison*Recall / (Precison + Recall)

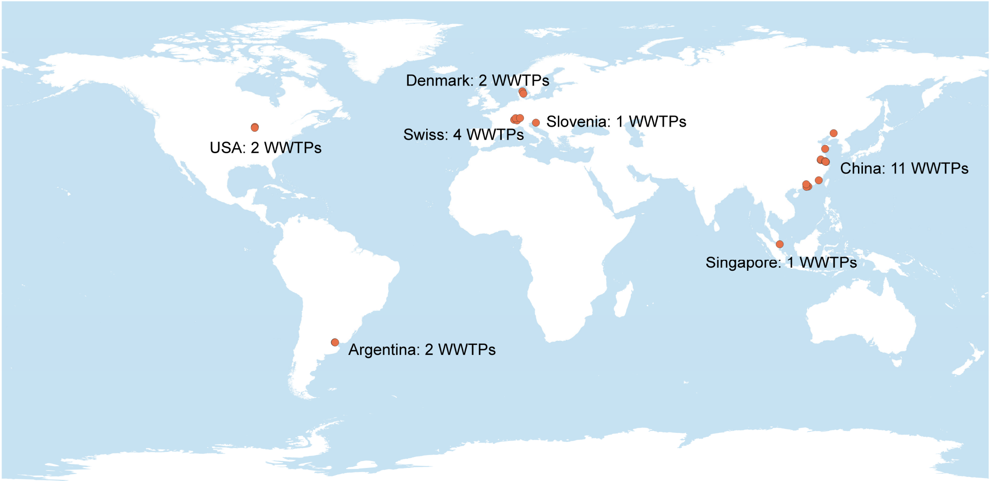


**Figure S1**. Geographical locations of the WWTPs where activated sludge samples were collected by us and other researchers.

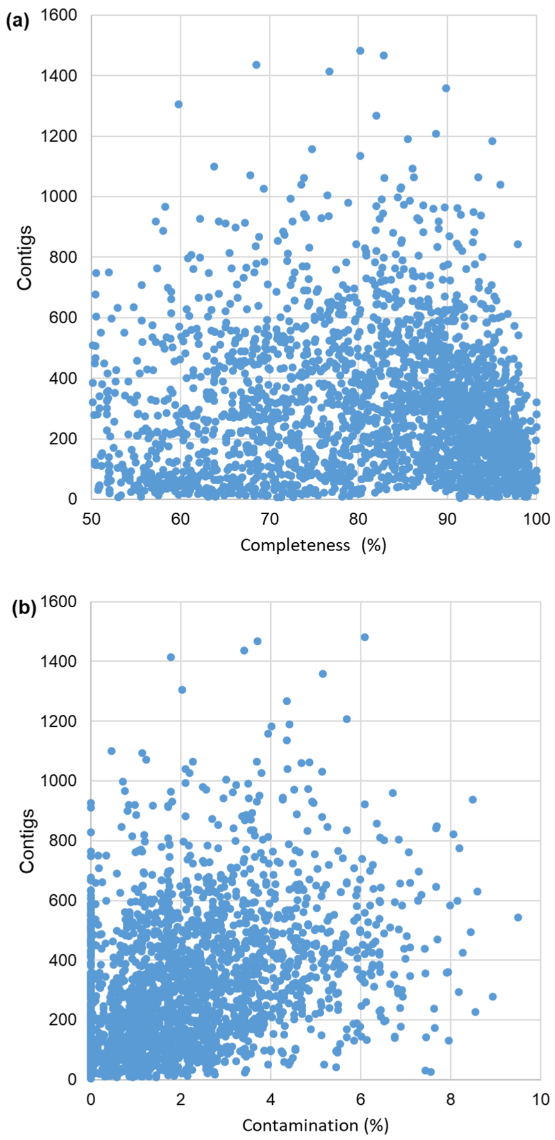


**Figure S2**. Associations between MAG completeness and number of contigs (a), and associations between MAG completeness and number of contigs (b).

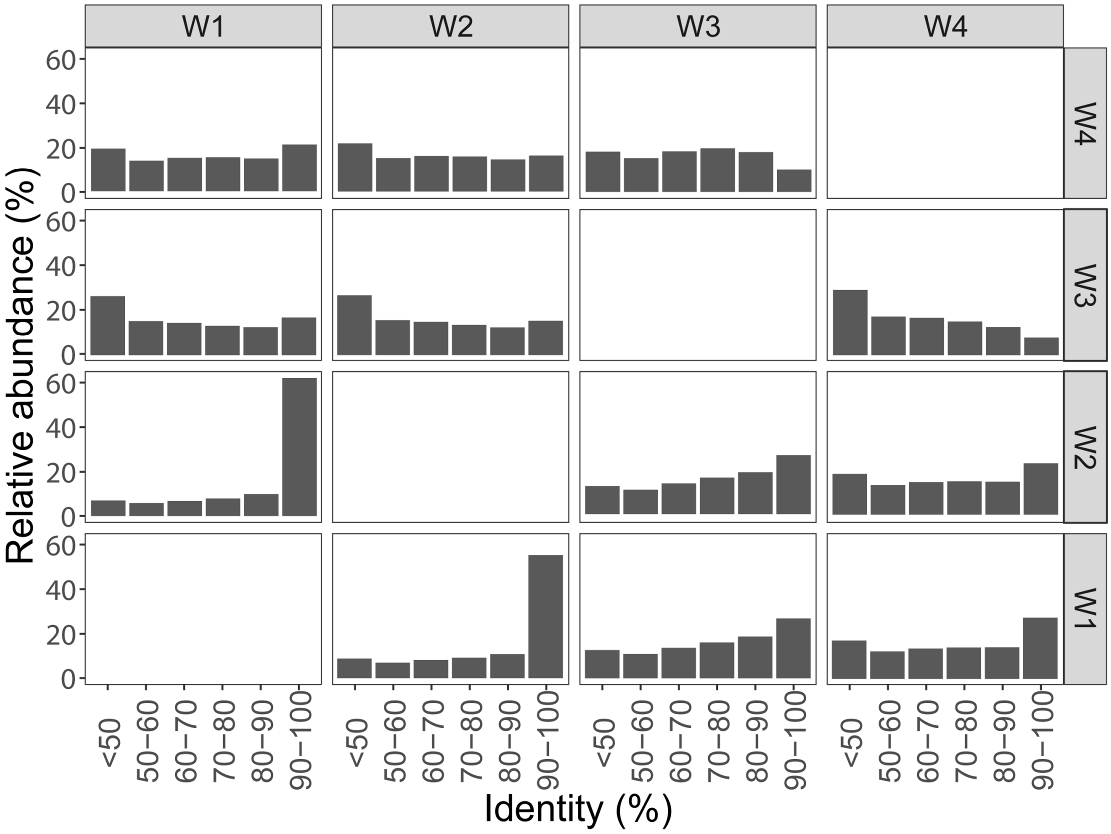


**Figure S3**. Profile of protein sequences identity between different WWTPs. The protein sequences predicted from all assembly contigs of each WWTP were compared each other with Diamond and then the best hits of the protein sequences were counted and summarized.

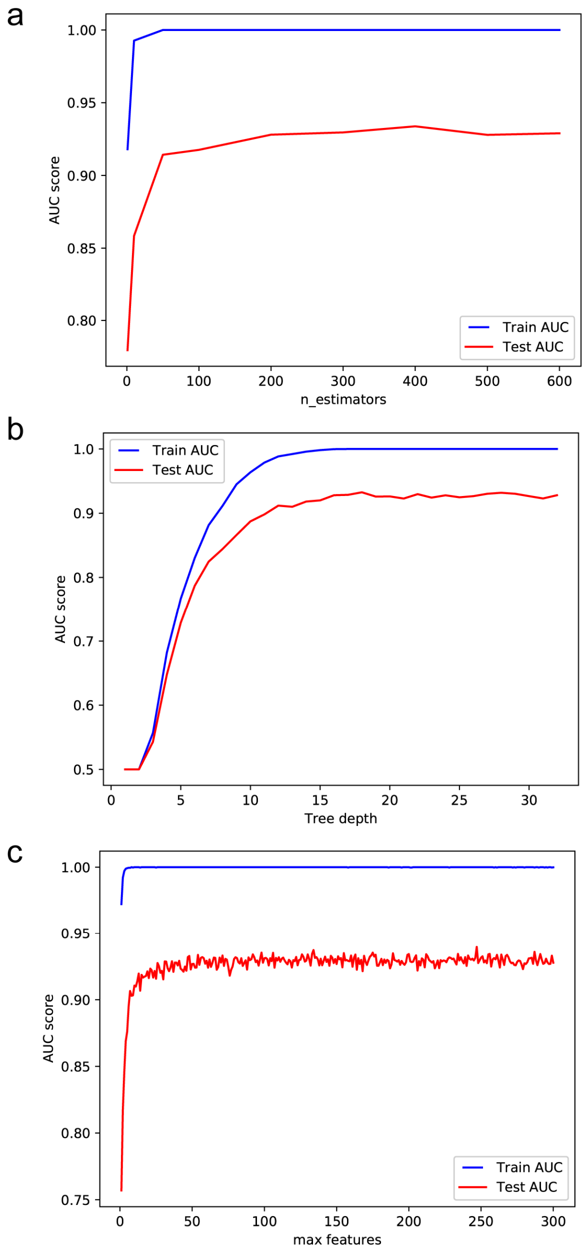


**Figure S4**. Random forest parameter tuning and optimization. (a) Number of trees (n_estimators); (b) Tree depth; (c) Maximum features.

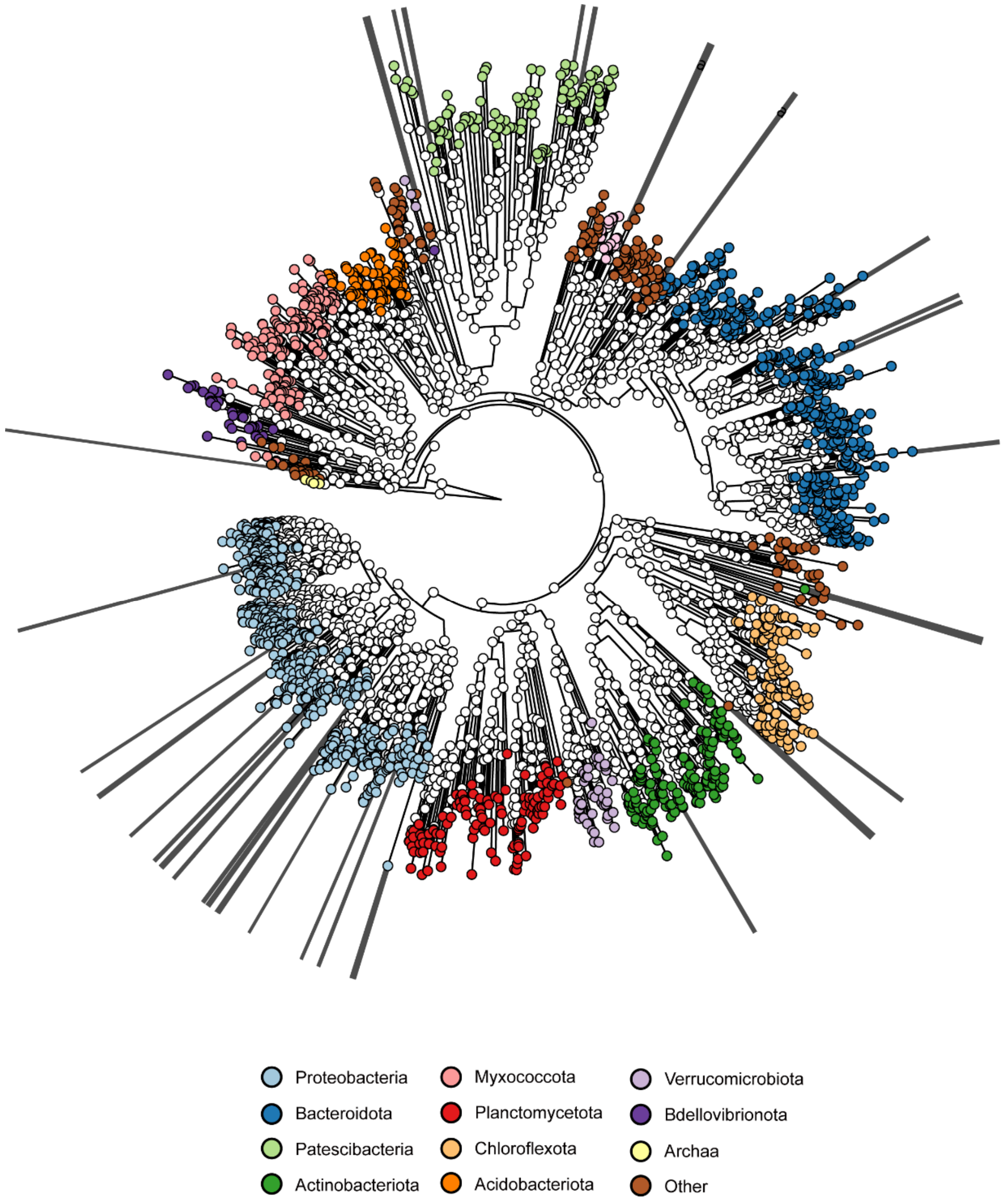


**Figure S5**. Phylogeny of the erroneously predicted MAGs. The topology of this tree is exactly same with Figure 1b. Extended lines were added to show positions of the erroneously predicted MAGs.
